# A generalizable cross-continent prediction of esophageal squamous cell carcinoma using the oral microbiome

**DOI:** 10.1101/2025.11.23.690048

**Authors:** Shahd ElNaggar, Wenlong Carl Chen, Leanne M. Prodehl, Thomas K. Marumo, Muhammed U. Khan, Christopher G. Mathew, Paul Ruff, Zhezhen Jin, Alfred I. Neugut, Anil K. Rustgi, Anne-Catrin Uhlemann, Tal Korem, Julian A. Abrams

## Abstract

Esophageal squamous cell carcinoma (ESCC) is a disease with limited tools for early screening and a poor prognosis. Symptoms typically appear late, and early cancer is hard to detect without endoscopic screening, which is inaccessible in most high-risk areas. Saliva is easily accessible, and its microbiome composition can serve as a marker for upper gastrointestinal tract disease. We studied the potential utility of an oral microbiome signature for ESCC in South Africa, a region with a high incidence of the disease. In a cohort of 48 ESCC patients and 110 controls, we found marked alterations in the oral microbiome in patients with ESCC, including significantly reduced alpha diversity and increased *Fusobacterium nucleatum*. We devised machine learning models that classify ESCC using microbiome data, finding good performance on held-out samples (area under receiver operating characteristic curve of 0.96), and demonstrated generalization to data across independent studies conducted in different geographic regions (0.64-0.81). Overall, our results demonstrate the potential of the oral microbiome to serve as a non-invasive screening tool for ESCC.

## Introduction

The dominant histological subtype of esophageal cancer globally is squamous cell carcinoma (ESCC), which has a very poor prognosis and remarkably high incidence rates in Eastern Asia and Eastern and Southern Africa, as opposed to esophageal adenocarcinoma (EAC), which predominates in Western countries^1^. Early detection of ESCC is a major clinical challenge. The lack of early symptoms and the inaccessibility of endoscopic screening in high-risk areas result in late-stage diagnoses and high mortality rates^2^. Established ESCC risk factors, including tobacco and alcohol use, polycyclic aromatic hydrocarbon exposure, hot food and beverage consumption, and poor oral health, can promote carcinogenesis^3,4^. However, these risk factors only partially explain the extremely high incidence of this disease in certain regions of the world and have not informed screening practices.

Increasing evidence suggests that the esophageal microbiome, which is in direct contact with the esophageal mucosa, may modulate the risk of epithelial cancers by way of immune activation and chronic inflammation^5^, and may influence ESCC treatment response^6,7^.

Additionally, previous studies have reported alterations of the esophageal and oral microbiome in ESCC^8,9^. Such associations may pave the way for the development of a microbiome-based biomarker for the detection of early-stage ESCC, which could potentially improve patient outcomes^10^. Our group previously showed that the salivary microbiome can distinguish patients with advanced precancerous changes and early esophageal adenocarcinoma^11^, suggesting that the salivary microbiome may also be useful for the identification of ESCC patients at early, treatable stages.

Given the known variation in the human microbiome across different populations and regions in the world, identifying robust associations between the microbiome and ESCC across various populations is essential for the development of a microbiome-based diagnostic. In this study, we focus on the oral microbiome of an understudied population in South Africa, where there is a high incidence of ESCC^12,13^. Several studies from China, which has regions of high ESCC incidence, have reported differences in the oral microbiome in ESCC patients^8,14–19^.

However, there remains a need to determine whether oral microbiome differences exist in other high-incidence regions of the world and whether they generalize across populations.

While direct sampling of the esophageal microbiome involves invasive procedures such as upper endoscopy, the sampling of saliva is relatively straightforward, and evidence shows that the salivary microbiome is strongly associated with the esophageal microbiome^20^. For this reason, the salivary microbiome may serve as a “window” to the esophageal microbiome and associated diseases. Additionally, salivary microbiome composition is temporally stable within individuals, suggesting its potential to provide diagnostic information regardless of sampling time^21,22.^

Here, we investigated whether the oral microbiome is a reliable marker of ESCC in a case-control study of 158 individuals from a high-incidence region in South Africa, including 48 ESCC patients and 110 matched controls. We found that the oral microbiome is significantly associated with ESCC and identified specific microbes, including *Fusobacterium nucleatum*, *Lautropia mirabilis*, *Veillonella dispar,* and *Prevotella salivae*, with higher abundance in individuals with ESCC. We then evaluated the generalizability of oral microbiome-based models in identifying ESCC patients across four studies, including ours and three others conducted in China, and showed that these models can identify individuals with ESCC from held-out studies. Overall, we found that compared to baseline clinical factors, the oral microbiome is an accurate predictor of ESCC.

## Results

### Patient characteristics from a South African ESCC case-control study

We enrolled 55 adult patients in Soweto, Johannesburg with histologically confirmed ESCC. We additionally enrolled 110 controls with no history of any cancer and no dysphagia symptoms.

At the time of sample collection, none of the patients had received treatment for ESCC, including chemotherapy, radiation, or surgery. We were unable to obtain histologic confirmation of ESCC for 7 cases and excluded these from analyses. Controls were geographically co-located, and all samples were processed together. We frequency matched controls to cases, to the extent possible, by age (±5 years), sex, and location, and characterized their salivary microbiomes using 16S rRNA gene amplicon sequencing (**Methods**). Patients with ESCC were older (two-sided Mann-Whitney U test p<0.001), and less likely to drink alcohol (chi-squared p=0.029), drink hot beverages (p=0.010), and cook outside (p=0.030; **Table S1**). There were no significant differences between ESCC and controls with regards to sex, marriage status, place of residence, smoking, self-reported HIV status, and diabetes status (**Table S1**).

### The oral microbiome in ESCC is distinguishable from controls

To investigate overall microbiome differences between ESCC patients and controls, we first tested whether within-sample diversity (Shannon α diversity) varied between groups. ESCC samples had significantly lower diversity (two-sided Mann-Whitney U test p=2x10^-4^; **Fig. 1a**). We observed no differences in α diversity based on age, sex, smoking, HIV status, or extraction batch (p>0.2 for all), although drinking hot beverages was associated with higher α diversity (p=0.027). Shannon diversity remained significantly associated with ESCC in a logistic regression adjusting for all available clinical and demographic covariates in addition to experimental batch (p=0.009). Additionally, we found that the oral microbiomes of ESCC patients clustered separately from those of controls when considering only the presence and absence of amplicon sequence variants (ASVs) (unweighted UniFrac, PERMANOVA p<0.0001; **Fig. 1b**), and this separation remained distinct even when adjusting for all available covariates (p<0.001). However, ESCC and controls did not cluster separately when microbiome distances were weighted by abundance (weighted UniFrac, p=0.096; **Fig. S1**). This suggests that rarer ASVs may be contributing to differences between ESCC and controls, as opposed to community-level differences.

**Figure 1.**
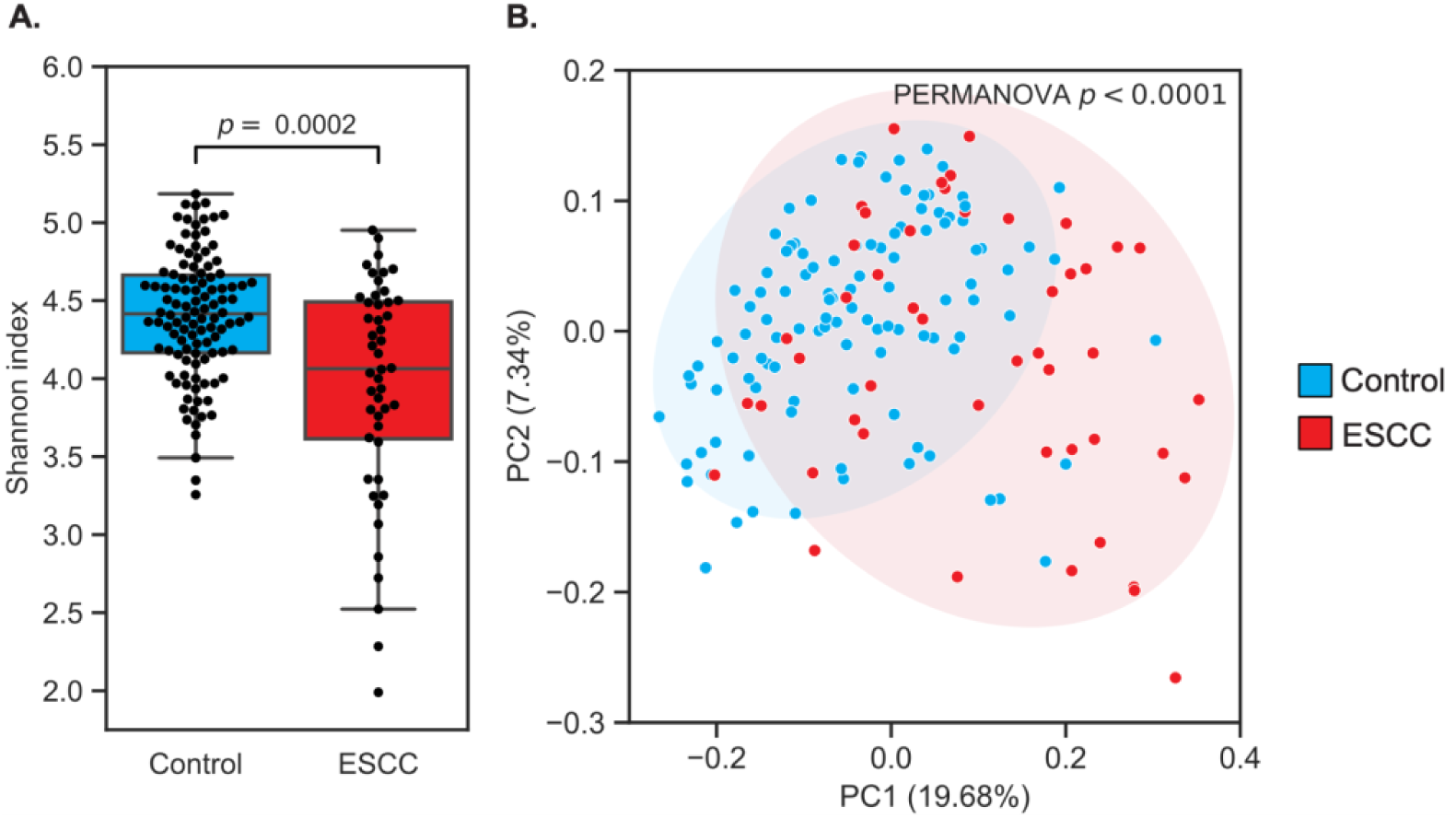
| The oral microbiome is associated with ESCC. **A.** Box and swarm plots (Box, IQR; line, median; whiskers, 1.5xIQR) showing significantly lower alpha diversity in ESCC patients compared to controls (two-sided Mann-Whitney U test p=2x10^-4^). **B.** PCoA of unweighted UniFrac distances demonstrated significant clustering of patients with ESCC (PERMANOVA p=10^-4^). Ellipses represent 2 standard deviations.

### Genus and ASV-level abundances are associated with ESCC

We next assessed whether specific genera were significantly associated with ESCC, with and without adjustment for covariates (**Methods**; see **Table S2** for unadjusted results). We found that *Capnocytophaga*, *Lautropia, Arachnia*, *Streptococcus, Selenomonas, Leptotrichia,* and *Campylobacter* were significantly elevated in the saliva of ESCC patients (FDR-corrected *P*<0.05 for all with covariate adjustment). Additionally, *Filifactor* and *Bacteroides* were depleted in the ESCC oral microbiome (FDR-corrected *P*=0.003 and *P*=0.01, respectively; **Fig. 2a**, **Table S2**).

**Figure 2.**
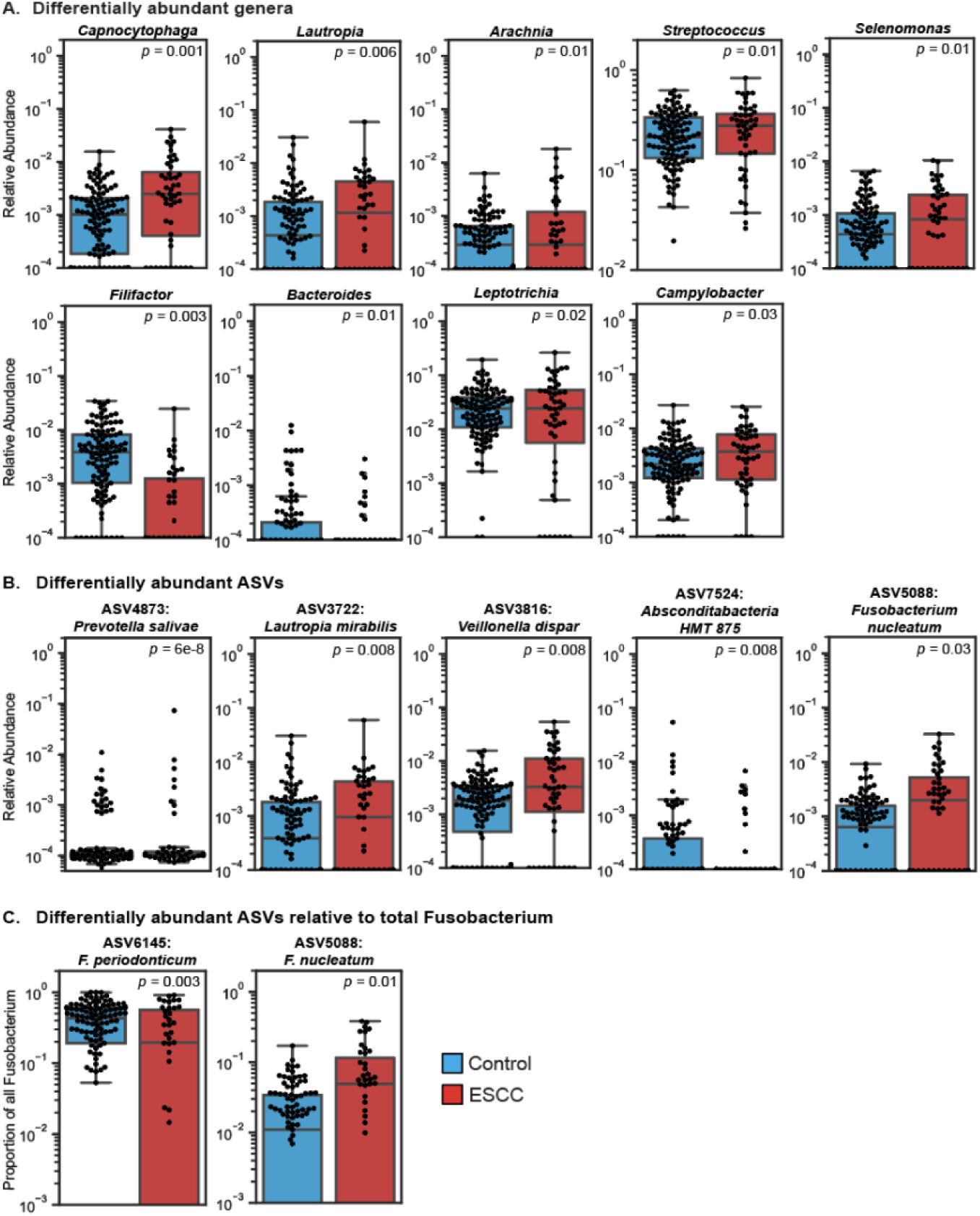
| Specific microbial taxa are associated with ESCC. A-C. Box and swarm plots (line, median; box, IQR; whiskers, nearest point to 1.5*IQR) showing the relative abundances (log scale) of differentially abundant genera (A), ASVs (B) and ASVs relative to the genus *Fusobacterium* (C) in the oral microbiome between ESCC and controls. Beta-binomial regression p values, calculated using corncob^62^, with FDR-correction (Benjamini-Hochberg) are displayed. Blue boxes represent controls (N=110) and red boxes represent ESCC cases (N=48).

Analyzing differential abundance of bacteria at the ASV level can offer more detailed insight than genus-level analyses, as microbial functions are often species-specific. Therefore, we investigated associations between ESCC and individual ASVs while adjusting for covariates (**Methods**). A total of five ASVs were identified as differentially abundant (FDR-corrected P < 0.05 for all): ASV4873 (*Prevotella salivae*), ASV3816 (*Veillonella dispar*), ASV3722 (*Lautropia mirabilis*), and ASV5088 (*Fusobacterium nucleatum*) were elevated in the ESCC oral microbiome, and ASV7524 (*Absconditabacteria* sp.) was depleted in ESCC (**Fig 2b**., **Table S3)**.

*F. nucleatum* is an oral commensal that is thought to promote the development of ESCC, is present in high abundance in a subset of ESCC cases, and is associated with worse clinical outcomes^23,24^. As we showed, higher abundance of oral ASV5088 (*F. nucleatum*) was associated with the ESCC oral microbiome. However, we did not identify genus-level *Fusobacterium* abundances associated with ESCC. We therefore next investigated whether certain ASVs were more likely to exist in higher proportions in the *Fusobacterium* genus between ESCC and controls, i.e. whether any ASVs were more prevalent relative to others within the genus. We found that ASV6145 (*Fusobacterium periodonticum*) was significantly decreased in ESCC relative to other *Fusobacterium* ASVs (p=0.003; adjusted for genus abundance in sample and ESCC risk factors). Additionally, ASV5088 (*F. nucleatum*) was significantly increased in ESCC relative to other *Fusobacterium* species (p*=*0.01) (**Fig 2c., Table S4)**. The dominance of *F. nucleatum* within the *Fusobacterium* genus in ESCC cases recapitulates its potential role in the disease.

### The oral microbiome accurately classifies ESCC

The ability of the oral microbiome to classify ESCC status holds great potential for the development of a non-invasive diagnostic. We therefore devised logistic regression models and checked whether the oral microbiome can be used to distinguish ESCC patients from controls. As a benchmark, we also tested models based on all available clinical information. Because the samples were processed in batches, we evaluated models on held-out batches (leave-one-batch-out cross-validation) to limit possible confounding effects^25,26^ (**Methods**). Model hyperparameters were tuned using the training set (“nested” cross-validation) without information leakage from the test set.

Models using clinical and demographic information obtained moderate accuracy with an area under the receiver operating characteristic curve (auROC) of 0.69 and an area under the precision-recall curve (auPR) of 0.54 (**Fig. 3a; Supplementary Fig. 3a)**. Using oral microbiome data at the ASV level, we were able to generate a model with significantly higher accuracy (auROC=0.96, auPR=0.92; DeLong’s test p=9.12 x 10^−9^ vs. model using clinical data). We also built a species-level model, which performed slightly worse compared to the ASV-level model (auROC=0.92, auPR=0.85; p=0.046 vs. ASV-level model). A model combining clinical and ASV-level oral microbiome data level did not improve upon the model based solely on oral microbiome data (auROC=0.96, auPR=0.93; p=0.72 vs. ASV-level model). This suggests that oral microbiome composition may be able to classify ESCC with high accuracy, and that its information content encompasses that of the associated clinical and demographic characteristics.

**Fig. 3.**
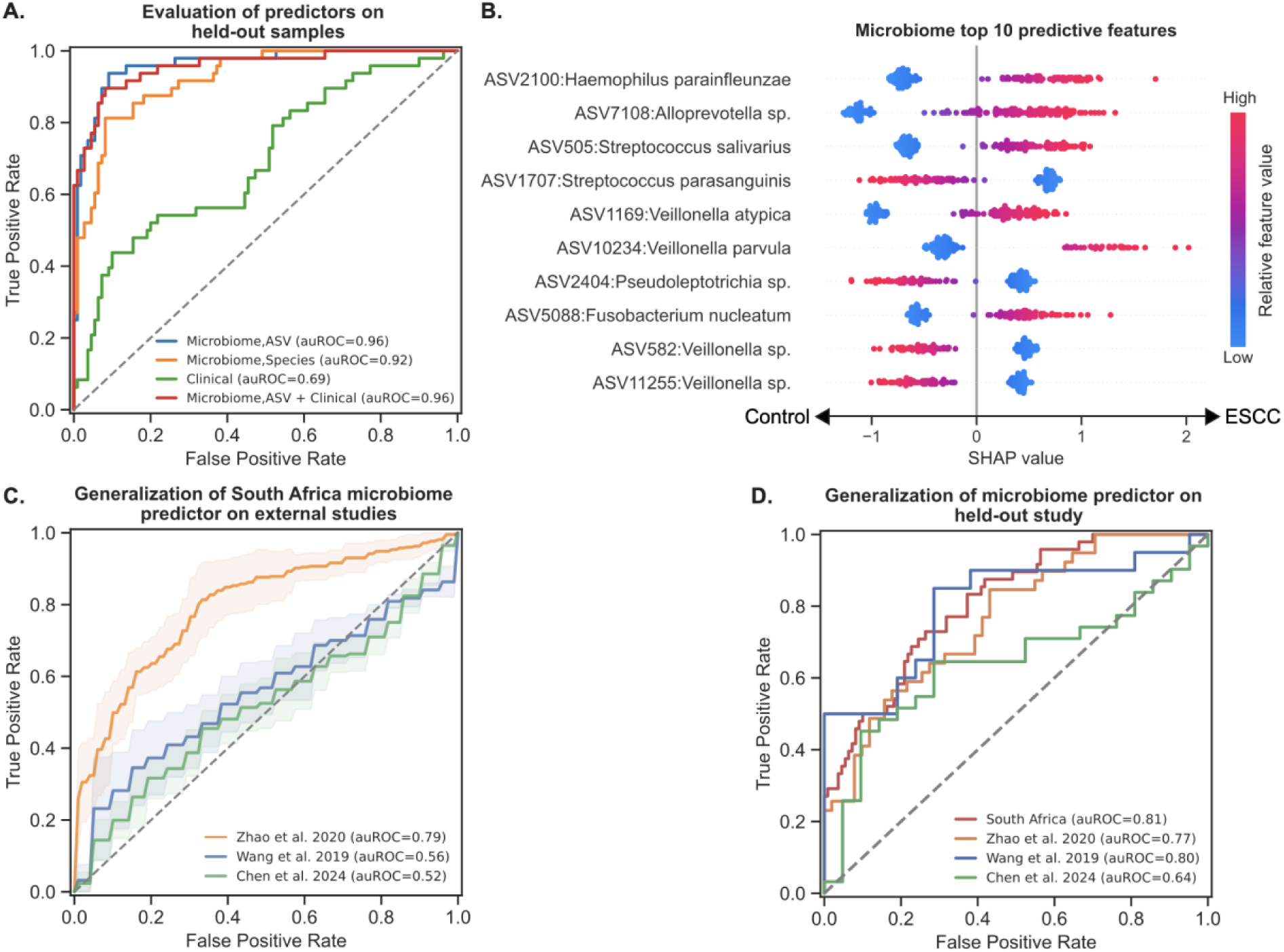
| Microbiome-based classification of ESCC. **A.** Receiver operating characteristic (ROC) curves comparing ESCC classification accuracy for ASV-level microbiome models (auROC=0.96),species-level models (auROC=0.92), models based on clinical data (auROC=0.69), and models based on both microbiome (ASV) and clinical data (auROC=0.96), evaluated on held-out experimental processing batches (**Methods**). **B.** Effect on prediction (SHAP values) for the top ten most predictive ASVs in the ASV-level microbiome-based model, sorted by importance. Each dot represents a specific sample, with the color corresponding to the relative value of the ASV in the sample compared to all other samples. **C.** ROC curves showing the performance of species-level microbiome predictors of ESCC, trained on our cohort (N=158) and evaluated separately on three held-out studies from China: Zhao et al. 2020 (N=91), Wang et al. 2019 (N=41), and Chen et al. 2024 (N = 52). The model from each external cross-validation fold was evaluated separately, with the line showing mean ROC curves and shaded regions representing ±1 standard deviation. **D.** ROC curves showing the performance of a species-level microbiome-based ESCC classifier, trained on all studies except one and evaluated on each held-out study.

We evaluated the importance of each feature towards the model prediction for each sample using SHapley Additive exPlanations (SHAP) values (**Fig. 3b** **and Fig. S2**). An analysis of our clinical-based predictor showed that age was one of the most predictive features, which corresponds to the older age of the ESCC group (**Supplementary Fig. 2a**). Additionally, of the top ten most predictive features identified in our microbiome-based predictor, ASV5088 (*F. nucleatum*) was the only taxon also highlighted as significantly differentially abundant in ESCC patients (**Fig. 2**). Interestingly, some taxa within the same genus had strongly opposing signals: for example, four species of *Veillonella* come up in model evaluation, two of which were associated with ESCC (ASV1169: *Veillonella atypica* and ASV10234: *Veillonella parvula*) and two other unidentified ASVs from this genus were associated with controls (ASV582 and ASV11255). Additionally, ASV505 (*Streptococcus salivarius*) was predictive of ESCC, while ASV1707 (*Streptococcus parasanguinis*) was associated with controls. These patterns indicate the importance of ASV-level analyses of oral microbiota in ESCC, as different microbes within a genus may play different roles in the oral cavity and in disease.

### Oral microbiome-based models for ESCC generalize across studies

Several studies have been performed in high-incidence regions of China assessing relationships between the oral microbiome and ESCC ^8,14–19,27–31^. Of the studies we identified, three included publicly available salivary microbiome 16S rRNA sequencing data from ESCC patients and non-ESCC controls: a study including 41 individuals (20 ESCC; Wang et al. 2019), all with periodontitis or gingivitis (gum disease)^15^; a study including 90 individuals (39 ESCC; Zhao et al. 2020)^18^; and a study including 52 individuals, 31 of which had early-stage ESCC and 21 controls^27^ (Chen et al. 2024). Wang et al. 2019 and Zhao et al. 2020 enrolled participants from the Henan, China region, albeit at different hospitals^15,18^, and Chen et al. 2024 of the studies enrolled participants from Nanjing, China^27^. All studies enrolled individuals who had not undergone previous treatment for ESCC, and their salivary microbiomes were profiled using the V3-V4 region of the 16S rRNA gene. Due to discrepancies in ASVs across these three studies and our study, which profiled the microbiome using the V1-V2 region, we were only able to evaluate our species-level microbiome-based model.

We first evaluated whether this model, trained on our cohort from South Africa, could independently generalize to each external cohort. We found that the model generalized well to Zhao et al. 2020 (auROC=0.79, auPR=0.77; p=2.87x10^-7^ one-sided Mann-Whitney U test; **Methods**), although not as well to Wang et al. 2019, which enrolled patients with gum disease (auROC=0.56, auPR=0.60; p=0.16), or to Chen et al. 2024, which focused on patients with early-stage ESCC (auROC=0.52, auPR=0.65; p=0.55; **Fig 3c, Supplementary Fig. 3b**). To assess whether this poor generalizability was due to less pronounced separation between ESCC and controls in the external studies, we ran 10-fold nested cross-validation separately on each external study (**Fig. S4**). We found that ESCC was highly distinguishable from controls using the species-level microbiome in Zhao et al. 2019^18^ (auROC=0.88, auPR=0.86; p=3.2x10^-10^) and was fairly distinguishable in the early-stage ESCC study^27^ (auROC=0.68, auPR=0.76; p=0.015). However, the oral microbiome was not able to distinguish ESCC well in the study consisting of only individuals with gum disease^15^ (auROC=0.54, auPR=0.50; p=0.33), which would explain the poor generalizability of our model to this particular study.

Finally, we evaluated the potential for a more global microbiome-based diagnostic by checking whether models trained on all studies except one could generalize to the held-out cohort. In this leave-one-study-out cross-validation, oral microbiome-based models were able to identify ESCC in the held-out study with some accuracy (auROC=0.64-0.81, auPR=0.70-0.84; all p<0.05; **Fig 3d, Supplementary Fig. 3c**), indicating a potentially generalizable microbiome signature for ESCC across geographic regions.

## Discussion

In this study, we characterized the oral microbiome of 158 individuals from South Africa using 16S rRNA gene sequencing, including 48 individuals diagnosed with ESCC and 110 healthy controls. We detected differences in the oral microbiome of ESCC compared to controls, including a significant decrease in microbial alpha diversity. We further found that genera including *Capnocytophaga, Lautropia*, *Arachnia*, *Streptococcus*, and *Selenomonas* were associated with ESCC, in addition to select ASVs including *F. nucleatum*, *V. dispar,* and *L. mirabilis*. Lastly, we demonstrated that microbiome-based models can classify ESCC status across geographically distinct cohorts, suggesting their potential as a screening tool for the disease. Prospective studies are necessary to evaluate whether the oral microbiome is unique prior to disease onset and if the microbiome can be used to classify the disease across cancer stages.

Many of the bacteria associated with ESCC identified in this study are known to promote inflammation and neoplasia in the oral cavity. *Capnocytophaga*, for example, has been demonstrated to invade oral squamous cell carcinoma cells and induce epithelial-to-mesenchymal transitions^32,33^. *Selenomonas* species, commonly associated with periodontal disease, are known to attach to gingival epithelial cells and trigger inflammatory responses^34,35^. *Streptococcus,* the most prevalent genus in the oral cavity, includes species such as *S. mitis* and *S. anginosus* that have been implicated in oral and upper gastrointestinal cancers^36–38^. A study from Tanzania analyzed the ESCC tumor-associated microbiome and reported similar findings compared to what we found in the oral microbiome, including high abundances of *Selenomonas, Streptococcus,* and *Campylobacter*^39^.

Our findings also support previous evidence implicating *F. nucleatum* in ESCC. *F. nucleatum* has been shown to be enriched in ESCC tumor tissue, and evidence suggests that the microbe may invade ESCC cells and enhance cell growth, thereby aiding disease progression^23,24,30^. Notably, we find that within the *Fusobacterium* genus, *F. nucleatum* tends to exist in higher proportions in ESCC compared to other species in the genera. By contrast, the oral microbiome of controls had a significantly higher proportion of *F. periodonticum*. Both *F. nucleatum* and *F. periodonticum* are known as active invader species, owing to their ability to independently invade epithelial host cells and, in the case of *F. nucleatum*, subvert host cell function^40–42^. Based on this evidence, it is possible that *F. nucleatum* could be outcompeting *F. periodonticum* in the ESCC oral microbiome.

Previous studies have established a higher risk of ESCC in those with poor oral health, specifically tooth loss and lack of regular oral hygiene^43–46^. Microbes involved in poor oral health may similarly be involved in inflammatory processes in ESCC, potentially explaining the link between poor oral health and cancer. A prospective study on esophageal adenocarcinoma (EAC) and ESCC found associations between *Treponema forsythia* and EAC as well as *Porphyromonas gingivalis* and ESCC^9^. These microbes are members of the “red complex” of periodontal pathogens, which are described as drivers of periodontitis^47^. In our study, we identified no relationship between ESCC and any red-complex species. Although we did not have oral health data for our participants, previous research suggests that these oral microbiome alterations may exist independent of oral health status. In a study on the oral microbiome in patients with Barrett’s esophagus and early EAC^11^, tooth loss was found to be associated with high-grade dysplasia and early EAC. However, even after adjusting for tooth loss, taxa associated with high-grade dysplasia and early EAC remained significant, indicating that the association of tooth loss with disease may be mediated through the oral microbiome.

The causal pathway between the oral microbiome, oral health, and ESCC risk remains undefined. A strength of our study is that we were able to devise accurate models for predicting ESCC using the oral microbiome in South Africa and across geographic regions, finding statistically significant signals even if not high accuracy (auROC of 0.64 for the least accurate model). Using cross-validation, we demonstrated that microbiome-based models were far superior to models using established clinical risk factors for the identification of patients with ESCC in South Africa. Additionally, we checked whether oral microbiome signatures of ESCC could generalize across diverse geographic regions. We evaluated species-level microbiome predictors on held-out studies, demonstrating that aggregate models generalized well to held-out studies, including one study that enrolled patients with early-stage ESCC. Lower performance of the individual South Africa-trained model on held-out studies from China can be explained by geographic effects, which are associated with the structure of the oral microbiome^48^, as well as discrepancies in extraction kits and sequencing protocols which may have introduced study-specific processing biases^49–51^. However, relatively good performance of a simple aggregate model across distinct cohorts, including a study with early-stage ESCC patients, demonstrates that there exists a somewhat generalizable oral microbiome signature that would allow for the identification of individuals with ESCC independent of geographic region and potentially in earlier stages of the disease. This is promising for the development of microbiome-based diagnostics. In order to further establish geographic generalization, models should be assessed in other areas with high incidence of ESCC such as Iran and East Africa^1^.

Additionally, while evidence suggests that the salivary microbiome is highly stable over time compared to other body sites^21^, future work should specifically establish whether a salivary microbiome-based diagnostic for ESCC remains accurate regardless of sampling time.

The study had certain limitations. Although staging data was not available for each participant, most ESCC patients presented with dysphagia, likely due to the presence of larger tumors and late-stage disease. The oral microbiome may be somewhat distinct in patients with early and late-stage ESCC, although our models showed some generalizability in the leave-one-study-out cross-validation setting between an external study with early-stage ESCC and other studies with later-stage ESCC, including our own. Since microbiome-based models trained on the South Africa study alone did not generalize as well to an external study including individuals with early-stage ESCC or another external study including only individuals with gum disease, further work must be done to ensure that a salivary microbiome-based diagnostic can distinguish ESCC at earlier stages and not be influenced by the presence of periodontal disease. Additionally, participants should have been matched more closely by age and excluded based on factors such as antibiotic use, although this was not possible due to the limited availability of participants. As controls did not undergo upper endoscopy, we cannot confirm with certainty that they did not have ESCC. However, this was a relatively healthy cohort without dysphagia symptoms, and thus the likelihood of undetected ESCC in out controls is very low.

Overall, these results demonstrate the potential of the oral microbiome to distinguish ESCC from controls and identify specific bacterial ASVs implicated in the disease. Translation of these findings for the oral microbiome-based screening of ESCC in low-resource settings will require validation across distinct geographic regions, in larger sample sizes, and especially in patients with early, potentially curable ESCC. Such a test could be used to triage patients for upper endoscopy. Additionally, modifying the oral microbiome in high-risk patients may also serve to lower the risk of ESCC, which could have a major public health impact in very high-incidence regions.

## Methods

### Study design

This cross-sectional study was designed to assess differences in oral microbiome composition between ESCC cases and matched controls. From September 2021 to November 2022, 55 Black South Africans with histologically confirmed ESCC were recruited across two GI clinics at the Chris Hani Baragwanath Academic Hospital in Soweto and the Charlotte Maxeke Johannesburg Academic Hospital in Johannesburg. Although patients did not undergo cancer staging, many individuals presented with dysphagia (trouble swallowing) and weight loss. Histological confirmation was obtained for all cancer cases. We excluded seven enrolled cases from our salivary microbiome analysis: histologic subtype could not be confirmed for 4 participants, one participant had EAC, one participant had gastric cancer, and one participant did not show evidence of cancer on histological assessment. Controls were enrolled 2:1 to ESCC cases from the Soweto area via PROMISE-SA, a multiple myeloma population-based screening study, between January 2022 and March 2023, and were frequency-matched to cases by age (+/- 5 years), sex, self-reported race, and study site. Matching by age was limited by the relatively small number of older eligible controls, resulting in a slight imbalance in age between cases and controls. It was confirmed via questionnaire that controls were between the ages of 40 and 75, self-identified as being Black or of African descent, had no current diagnosis of any cancer, and had no symptoms of dysphagia. Neither cases nor controls had a previous diagnosis of head or neck cancer, or any other type of cancer. At the time of saliva collection, none of the patients had received treatment for ESCC, including chemotherapy, radiation, or surgery.

We collected demographic and clinical data from each participant. Additionally, we collected 2 mL of saliva using the Oragene OG-500 DNA Saliva Collection Kit (DNAGenotek, Ontario, Canada). In our microbiome analyses and predictive modeling, we included the following clinical metadata variables due to their potential relevance to ESCC and/or salivary microbiome composition: patient age (years), sex, marriage status, location of residence, cooking location, highest level of education completed, smoking, alcohol consumption, hot beverage consumption HIV status, and history of diabetes mellitus. Clinical and demographic data were unavailable for two controls. Additional data is missing for age (3 ESCC), sex (2 ESCC), location of residence (2 ESCC; 1 control), cooking location (3 ESCC), education (3 ESCC), smoking (2 ESCC), alcohol consumption (2 ESCC; 1 control), hot beverage consumption (5 ESCC), diabetes status (2 ESCC), and HIV status (4 ESCC; 21 controls). In Supplementary Table 1 only, education levels Grade 1 through 7 are collapsed into “Primary school” and Grade 8 through 12 are collapsed into “Secondary school.”

All enrolled patients provided written consent. The study was conducted in accordance with the principles of the Declaration of Helsinki and approved by the Human Research Ethics Committee at the University of the Witwatersrand (certificate number: M180306) and by the Institutional Review Board at Columbia University.

### Statistics and Reproducibility

Sample size calculations were made based on differences in α diversity between cases and controls; a projected sample size of 50 cases and 100 controls provided 82% power to detect a 0.5 standard deviation difference, assuming a type I error rate of 0.05. Between-group comparisons of demographic variables in **Table S1** were performed using chi-squared tests for categorical variables and Mann-Whitney U tests for continuous variables. Unadjusted ⍺-diversity comparisons between cases and controls were conducted using Mann-Whitney U tests, and adjusted comparisons were conducted using a logistic regression model that included the clinical and demographic variables listed above. β-diversity comparisons were conducted using permutational multivariate analysis of variance (PERMANOVA) with 10,000 permutations to calculate p values. Adjusted PERMANOVA tests included the clinical and demographic variables listed above in the model. The DeLong test was used to assess statistically significance of differences in machine learning model performance and Mann-Whitney U tests were used to assess whether models performed better than a random classifier. Statistical significance was defined as p<0.05. In the case of multiple comparisons, reported p values are corrected for false discovery rates (FDR) using the Benjamini-Hochberg method^63^.

### Microbiome sequencing and analysis

Microbial DNA was isolated in 11 separate batches from saliva specimens using the QIAamp BiOstic Bacteremia DNA kit as per the manufacturer’s protocol. Negative extraction blanks were processed in tandem for 9 out of 11 batches. The V1-V2 region of the 16S rRNA gene was amplified using Illumina adapter-ligated primers (27F-338R). The resulting libraries were barcoded and sequenced using the Illumina MiSeq platform.

To limit the analysis to bacterial reads only, reads mapped with bowtie2 (ref. ^52^) with inclusive parameters to CHM13-V2 human and PhiX genome sequences were removed from further analysis. The QIIME 2 v2024.2 microbiome analysis platform, which is a wrapper for various microbiome bioinformatics tools, was used for the following analyses^53^. DADA2 v1.26.0 was used to filter low-quality reads, merge paired-end reads, and to identify all amplicon sequence variants (ASV) and their counts in each sample^54^. A median of 44,788 clean and non-chimeric paired-end reads were available per sample for analysis. All ASVs were aligned with mafft^55^ and used to construct a phylogeny with fasttree2^56^. Taxonomies were assigned to ASVs using the q2-feature-classifier classify-sklearn naive Bayes taxonomy classifier^57^. A classifier trained on Human Oral Microbiome Database v15.23 (HOMD) OTU sequences was used to assign taxonomies to each ASV^58^, except in cross-study species-level analyses where we used Greengenes2 2022.10 (detailed below).

Possible bacterial contaminants were removed *in silico* using SCRuB, which probabilistically estimates the true counts in a sample based on the proportions of taxa observed in the negative extraction blanks^59^. Nine out of eleven batches were “decontaminated” using their corresponding blanks, while the two batches without corresponding blanks were decontaminated using a pooled blank which summed taxa across all available blanks.

Sample α diversity was estimated using the Shannon index, and inter-sample diversity (β diversity) was estimated using unweighted and weighted Unifrac^60^. Prior to diversity estimates, rare taxa were filtered (only taxa present in at least 5 samples with a total count of at least 20 were included) and each sample was rarefied to 9,700 reads, which was the minimum read depth across samples. 2,091 out of 11,277 ASVs were retained post-filtering. Community-level differences between ESCC samples and control samples were tested via Permutational multivariate analysis of variance (PERMANOVA) using the “adonis” function within the QIIME2 q2-diversity plugin, once without adjusting for covariates and once adjusting for all available clinical and demographic covariates, as well as batch^61^.

### Differential abundance testing

Differential abundance analysis was performed in R v4.3.2 using beta-binomial regression models from Corncob v.4.1.^62^. ASV tables were filtered to include taxa present in five or more samples. Models were adjusted for batch, age, alcohol consumption, cooking location, hot beverage consumption, and smoking. To identify differentially abundant taxa, we first assessed which taxa had an FDR-corrected p-value of less than 0.1 in a model that is only adjusted for batch and then identified which of these taxa were also significant (FDR-corrected p<0.05) in the full model, which included the rest of the covariates. For the *Fusobacterium* dominance analysis, models were additionally adjusted for the total abundance of the genus in each sample.

Reported p-values are corrected for false discovery rate (FDR) using the Benjamini-Hochberg method^63^; taxa with significant p-values are displayed in **Tables S2** (genus), **S3** (ASV), and **S4** (*Fusobacterium* ASVs).

### Training, testing, and evaluation of ESCC classifiers

Supervised prediction models were built to classify ESCC versus control samples in our cohort using the scikit-learn Python library (v1.3.1). Logistic regression was used for all classification tasks. Model performance was evaluated across four feature sets: (1) clinical and demographic variables (including age, sex, marriage status, education, residence location, cooking location, smoking, alcohol consumption, hot beverage consumption, HIV status, and diabetes status); (2) microbiome data at the ASV level (11,277 features); (3) microbiome data at the species level (Greengenes2; 576 features); and (4) a combined dataset consisting of both clinical and microbiome (ASV) data (i.e., 1+2).

For all models, samples were split up into training and test sets by grouping samples by batch and creating eleven test-train splits, leaving one batch as a held-out test set each time (i.e. eleven-fold cross validation). This was done to account for confounding effects in our models introduced by class imbalances in each batch (e.g. some batches consisted only of ESCC samples). For the clinical models, missing information was imputed using the median of the training set for continuous variables and the mode of the training set for categorical variables. To tune hyperparameters, we used nested cross-validation. Model hyperparameters in our microbiome models included steps for pre-processing and feature selection. All features with a variance of 0 were removed and the remaining features were center log-ratio (CLR) transformed. Feature selection was implemented as part of the nested cross-validation procedure to ensure that feature selection was performed only on the training data in each fold, thereby preventing information leakage in the validation or test set. We fit a lasso model to the data and then retained all features with non-zero coefficients. We tuned a hyperparameter *k*, which controls the proportion of features to retain. For each value of *k*, we determined the optimal L1 regularization strength (α) that results in approximately *k* percent of the input features having nonzero coefficients. Each training set was split into five folds (i.e. ‘inner folds’) on which we used 1,000 iterations of a random set of hyperparameters. This process was repeated five times to account for stochasticity. The best hyperparameter set was selected as the model with the highest average area under the receiver operating characteristic curve (auROC) score based on performance on the inner folds. This model was then trained on the entire training data and then evaluated once on the unseen held-out batch. For model evaluation, we calculated the overall auROC and area under the precision recall curve (auPR) across all folds together.

### Cross-study preprocessing and validation of ESCC classifiers

We conducted a literature search to identify studies suitable for external validation of our microbiome-based ESCC predictor. 12 studies were initially identified that sampled the oral and/or esophageal microbiome of individuals with ESCC and non-ESCC controls^8,14–19,27–31^. We excluded studies without publicly available 16S rRNA sequencing data of saliva samples. We reprocessed PRJNA660092^18^, PRJNA587078^15^, and PRJNA961904^27^ to the ASV-level using the same steps detailed above. For all three cohorts, including our own, we assigned taxonomies using the q2-feature-classifier classify-sklearn naive Bayes taxonomy classifier trained on the Greengenes2 2022.10 database, with the ‘–p-confidence’ parameter set to 0. This approach ensured species-level assignments for all ASVs, regardless of confidence scores, to enable a consistent species-level comparison across all studies. We filtered out features that appeared in less than 10% of samples, which resulted in 315 features being retained, and CLR-transformed the remaining features. Then, we performed leave-one-study-out cross-validation, where we trained a logistic regression with no penalty on all but one study, and evaluated the model’s performance on each held-out study. For the species-level model trained on our cohort and tested on the external cohorts, we retrained each nested model on the entire South Africa cohort and individually tested each of the eleven models on the external cohorts. Lastly, we performed 10-fold nested cross-validation within each external study using species-level microbiome data. Hyperparameter optimization and preprocessing steps on the external studies were identical to those used on our data. To assess whether models performed better than a random classifier, we conducted a one-sided Mann-Whitney U test to evaluate whether predicted scores were higher for the positive than for the negative class.

## Acknowledgments

This study was supported by NIH/NCI R01 CA238433-S1. Additional support was provided by the Herbert Irving Comprehensive Cancer Center (NIH/NCI P30CA013696-50), the South African Medical Research Council (Extramural – CECRC), and the South African National Research Foundation (Thuthuka). The project on which this publication is based was in part funded by the German Federal Ministry of Education and Research 01KA2220B. This research was funded in part by the Science for Africa Foundation to the Developing Excellence in Leadership, Training and Science in Africa (DELTAS Africa) program [Del-22-008] with support from Wellcome Trust and the UK Foreign, Commonwealth & Development Office and is part of the EDCPT2 programme supported by the European Union. We thank the CUIMC Microbiome & Pathogen Genomics Core for 16S rRNA sequencing, and members of the Korem group for useful discussions.

## Author contributions

J.A.A., W.C.C., C.G.M., A.I.N., and Z.J. conceived and designed the study. Recruitment and clinical assessment of participants were performed by W.C.C., L.M.P., T.K.M., and M.U.K. A.C.U. generated all data. S.E., Z.J., T.K., and J.A.A designed all analyses, conducted data analysis, and interpreted the results. All authors (S.E., W.C.C., L.M.P., T.K.M., M.U.K., C.G.M., P.R., Z.J., A.I.N., A.K.R., A.C.U., T.K., J.A.A.) contributed to writing and editing the manuscript and approved the final version. J.A.A. and T.K. supervised the study.

---

**Supplementary Figure 1.**
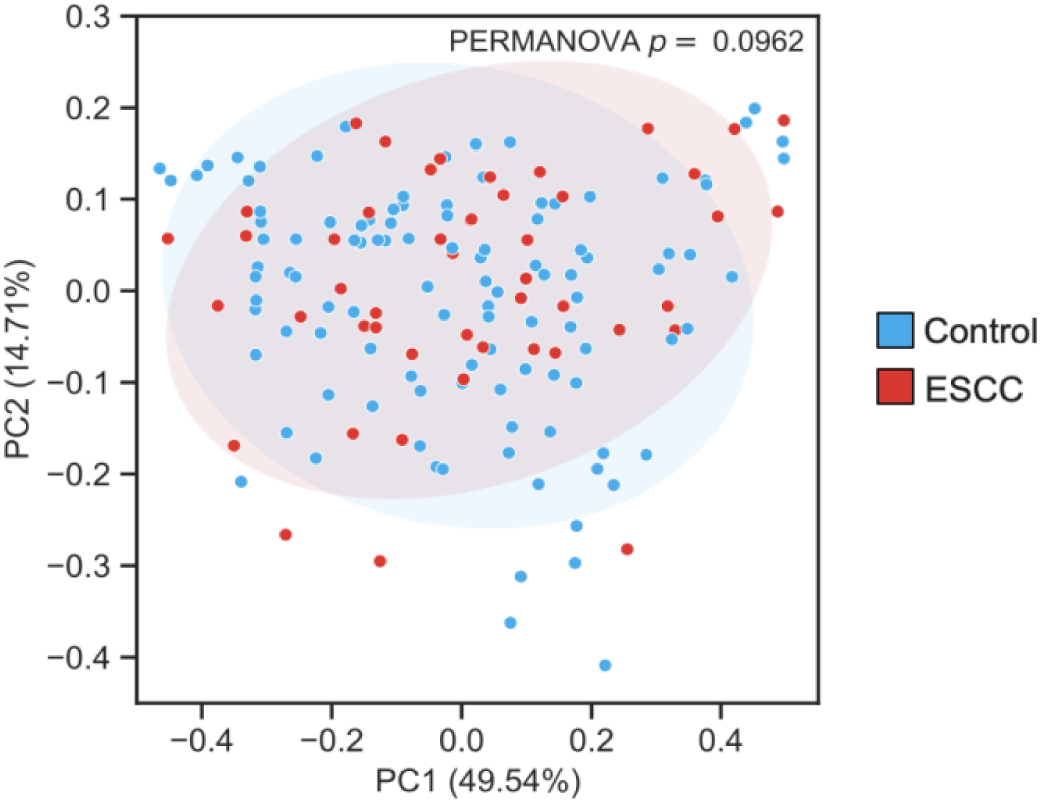
| Weighted UniFrac PCoA demonstrated no association with ESCC. PCoA on Weighted UniFrac distances demonstrated no significant clustering of patients with ESCC (PERMANOVA P = 0.096). Ellipses represent 2 standard deviations.

**Supplementary Figure 2.**
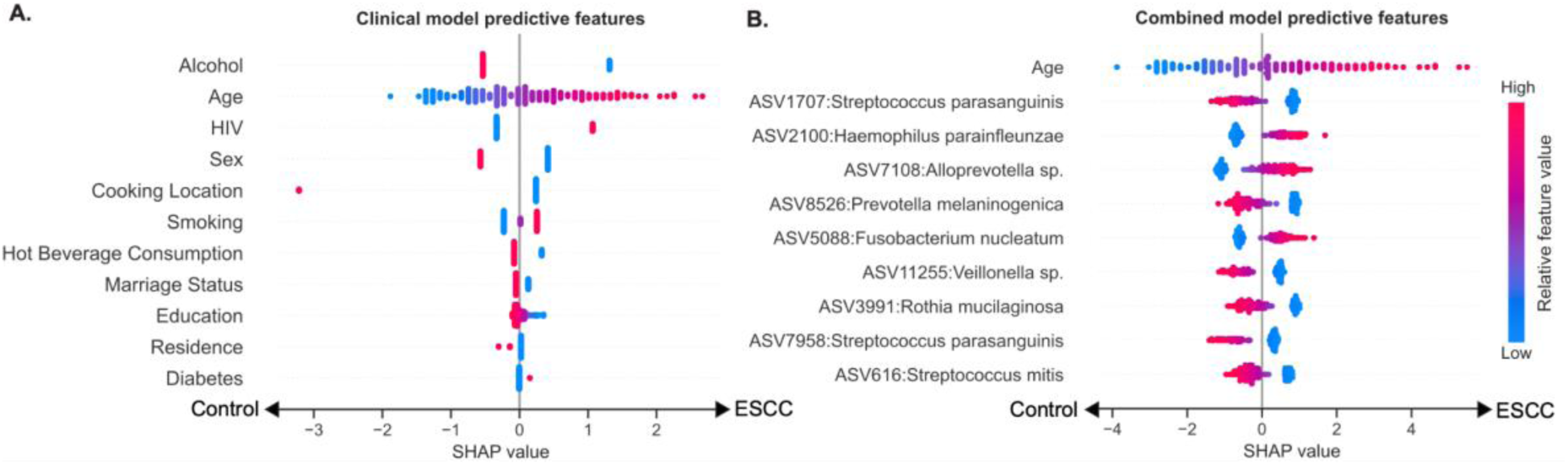
| Top predictive features in the clinical and combined model. Effect on prediction (SHAP values) for the topmost predictive covariates in the **A.** clinical model and the **B.** combined clinical and microbiome-based model, with features sorted by importance. Each dot in the plots represents a specific sample, with the color corresponding to the value of the feature in the sample compared to all other samples.

**Supplementary Figure 3.**
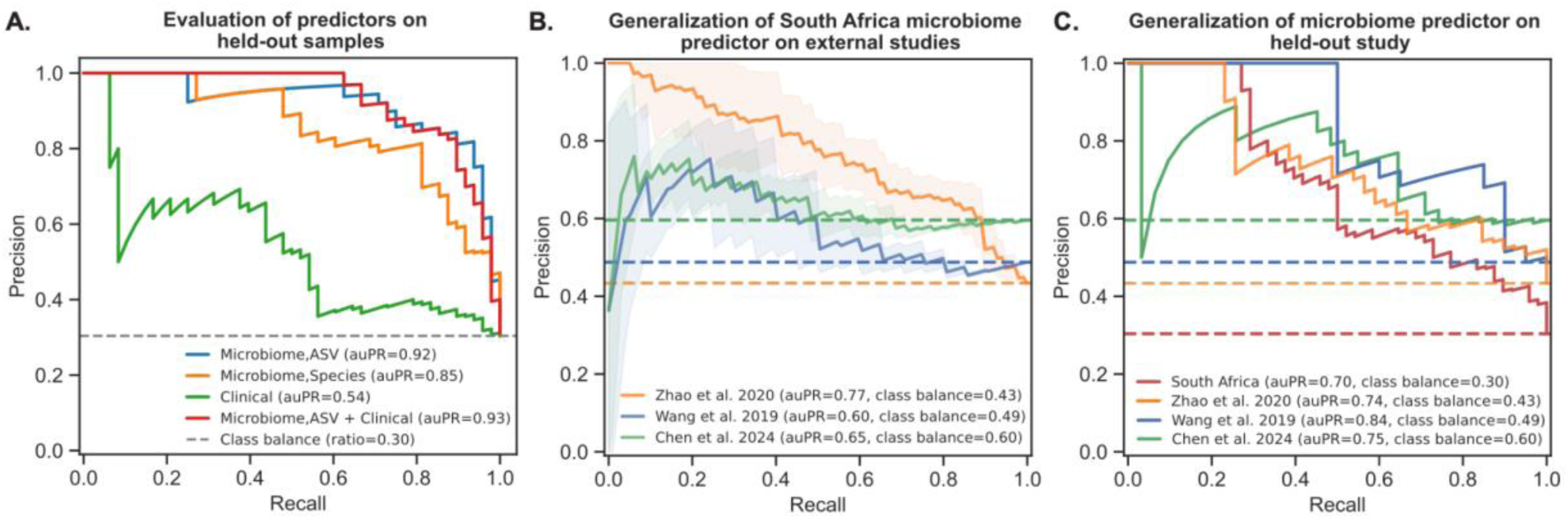
| Microbiome-based classification of ESCC **A.** Precision-recall (PR) curves showing the classification accuracy for ASV-level microbiome-based models (auPR=0.92),species-level models (auPR=0.85), models based on clinical data (auPR=0.54), and models based on both microbiome (ASV) and clinical data (auPR=0.93), all evaluated on held-out experimental processing batches. **B.** PR curves showing the classification accuracy of species-level microbiome predictors of ESCC, trained on our cohort (N=158) and evaluated separately on held-out studies from China: Zhao et al. 2020 (N=91), Wang et al. 2019 (N=41), and Chen et al. 2024 (N = 52). The model from each external cross-validation fold was evaluated separately, with the line showing the mean PR curve and shaded regions representing ±1 standard deviation. **C.** PR curves showing the performance of a species-level microbiome-based ESCC classifier, evaluated on each held-out study. The class balance for each study is shown as a dashed horizontal line.

**Supplementary Figure 4.**
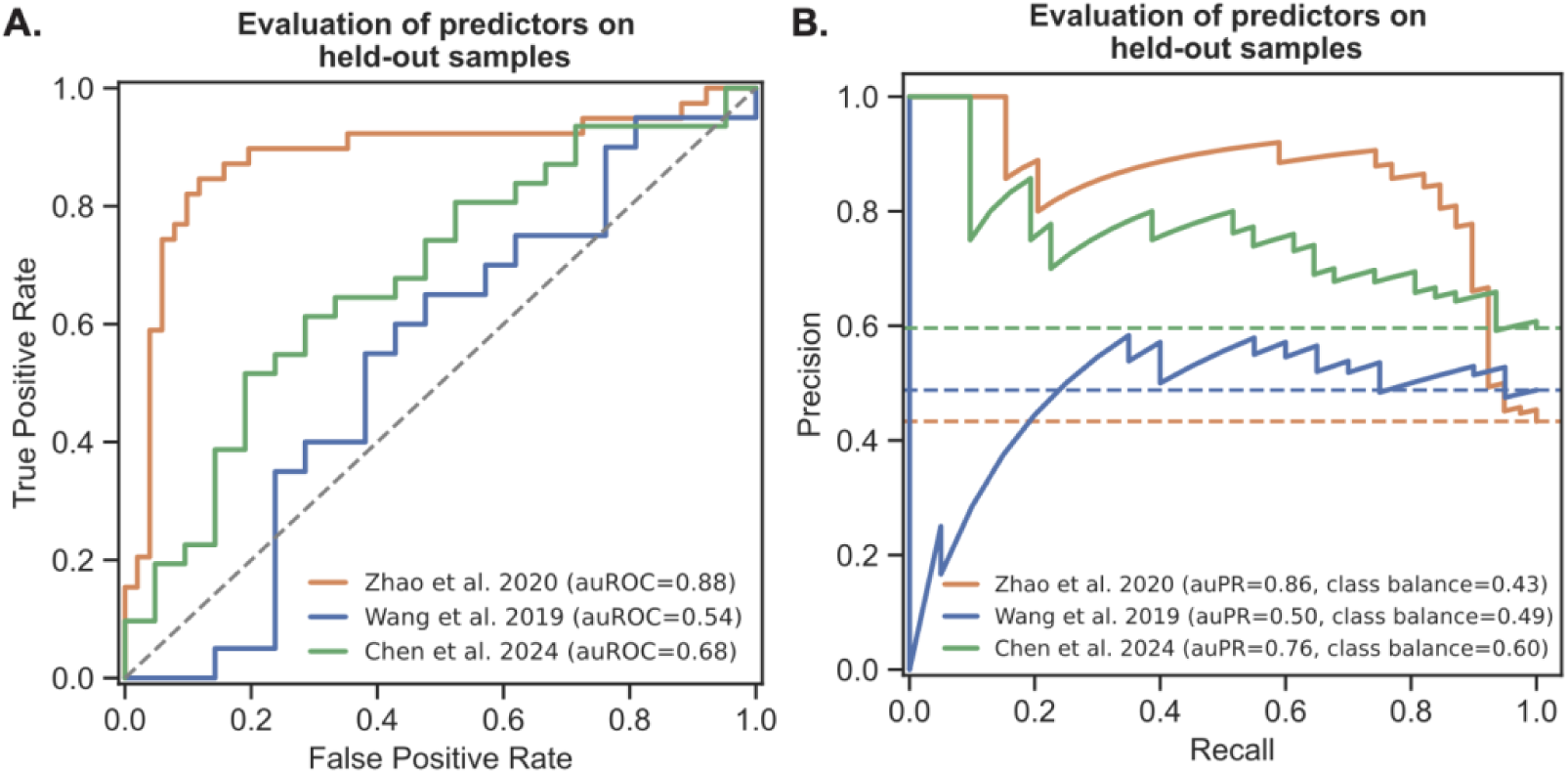
| Microbiome-based classification of ESCC in external studies A,B. Receiver operating characteristic (ROC) curves (A) and precision-recall (PR) curves (B) showing the classification accuracy of species-level microbiome-based models, evaluated on held-out samples using 10-fold nested cross-validation within each external study for Zhao et al. 2020, Wang et al. 2019, and Chen et al. 2024.

**Supplementary Table 1.**
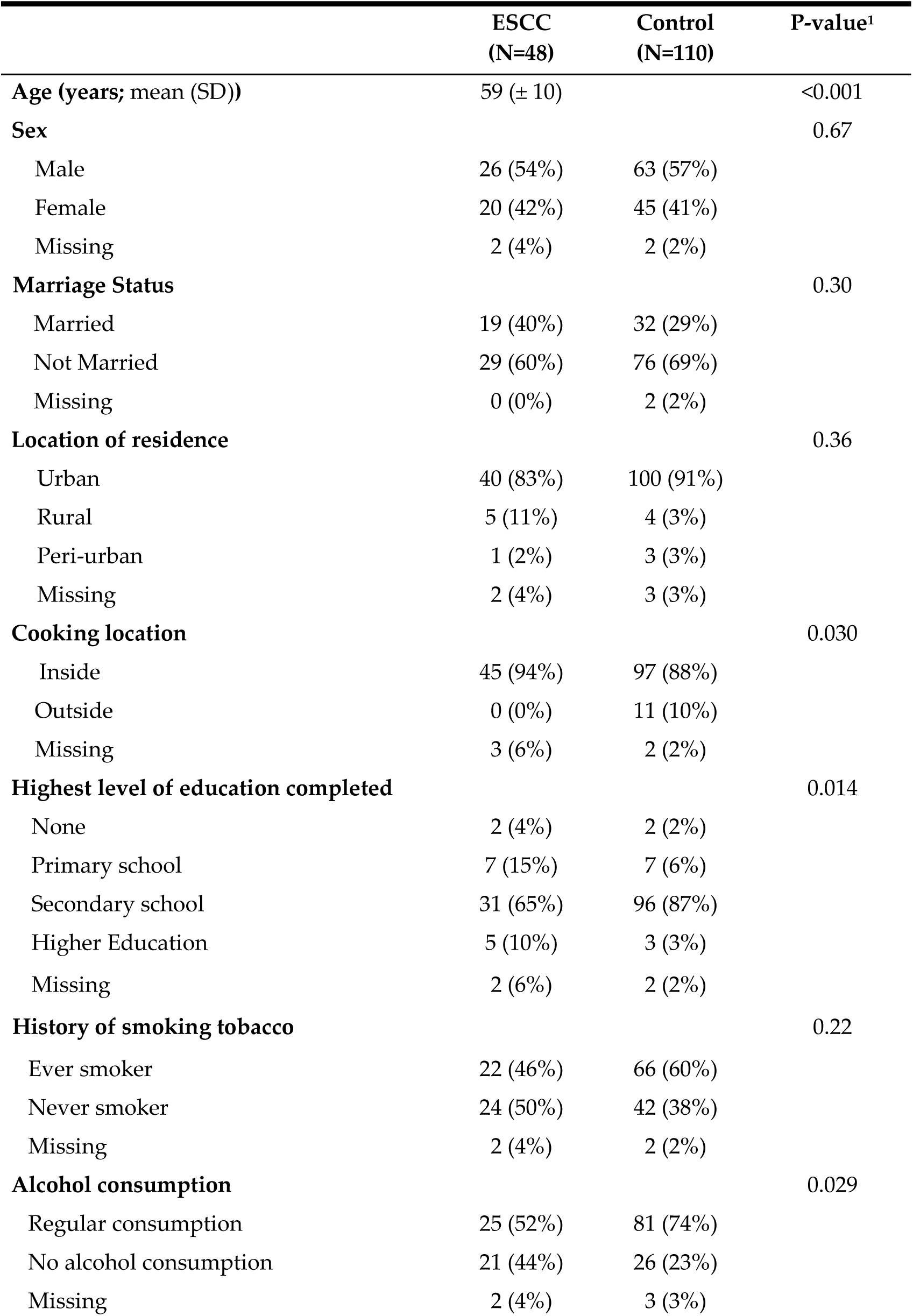
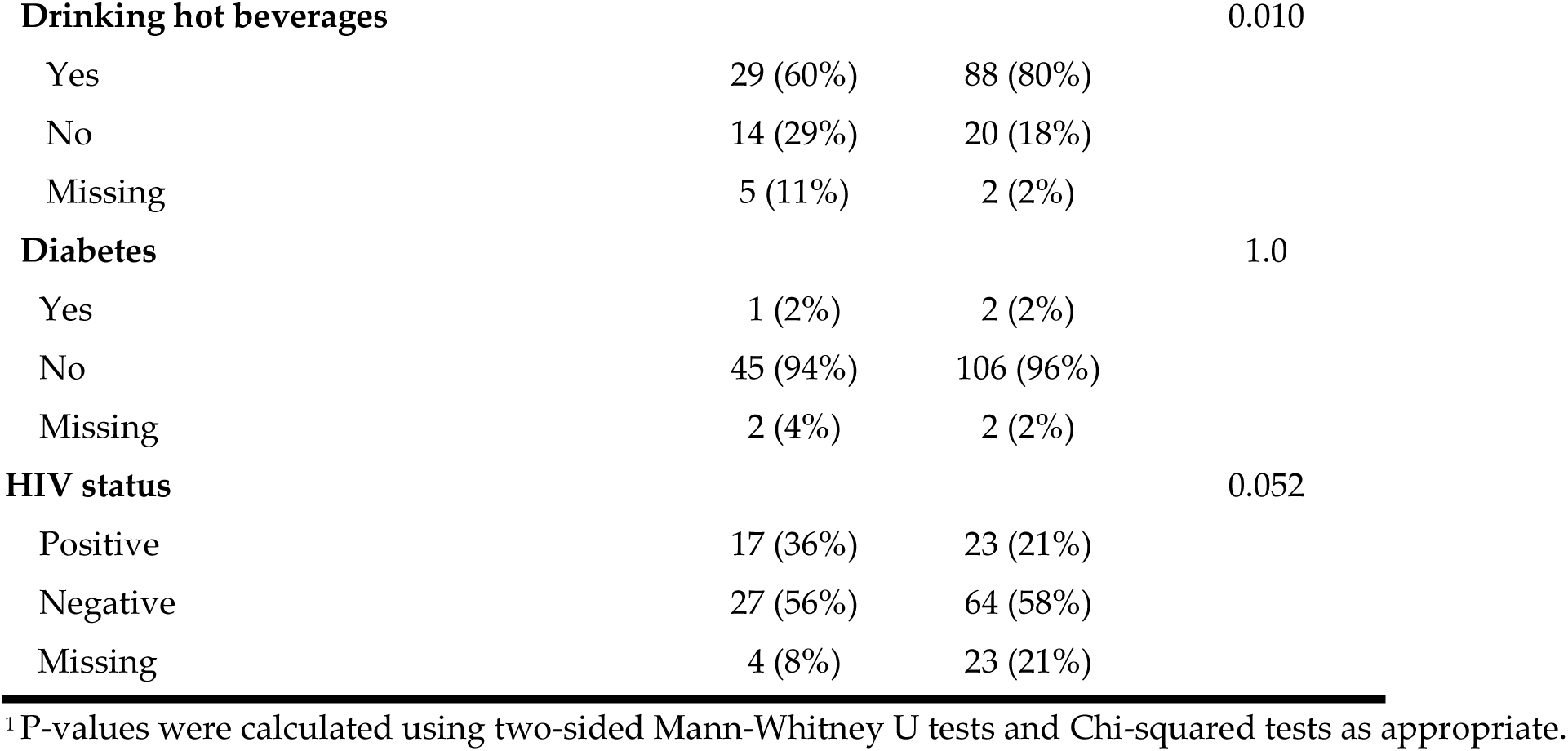
Patient characteristics

## The following tables are included as a supplementary file

Table S2 | Genus differential abundance unadjusted and adjusted beta-binomial regression results

Table S3 | ASV differential abundance unadjusted and adjusted beta-binomial regression results

Table S4 | *Fusobacterium* ASV differential abundance unadjusted and adjusted beta-binomial regression results

## References

1. Bray, F. et al. Global cancer statistics 2022: GLOBOCAN estimates of incidence and mortality worldwide for 36 cancers in 185 countries. CA Cancer J. Clin. 74, 229–263 (2024).

2. Codipilly, D. C. et al. Screening for esophageal squamous cell carcinoma: recent advances. Gastrointest. Endosc. 88, 413–426 (2018).

3. Chiang, H.-C., Hughes, M. & Chang, W.-L. The role of microbiota in esophageal squamous cell carcinoma: A review of the literature. Thorac Cancer 14, 2821–2829 (2023).

4. Ohashi, S. et al. Recent Advances From Basic and Clinical Studies of Esophageal Squamous Cell Carcinoma. Gastroenterology 149, 1700–1715 (2015).

5. Sharma, T. et al. Cross-talk between the microbiome and chronic inflammation in esophageal cancer: potential driver of oncogenesis. Cancer Metastasis Rev. 41, 281–299 (2022).

6. Wu, H. et al. Intratumoral Microbiota Composition Regulates Chemoimmunotherapy Response in Esophageal Squamous Cell Carcinoma. Cancer Res. 83, 3131–3144 (2023).

7. He, Y., Li, X.-Y., Hu, A.-Q. & Qian, D. Salivary microbiome is associated with the response to chemoradiotherapy in initially inoperable patients with esophageal squamous cell carcinoma. J Oral Microbiol 16, 2359887 (2024).

8. Chen, X. et al. Oral Microbiota and Risk for Esophageal Squamous Cell Carcinoma in a High-Risk Area of China. PLoS One 10, e0143603 (2015).

9. Peters, B. A. et al. Oral Microbiome Composition Reflects Prospective Risk for Esophageal Cancers. Cancer Res. 77, 6777–6787 (2017).

10. Wang, G.-Q. et al. Long-term results of operation for 420 patients with early squamous cell esophageal carcinoma discovered by screening. Ann. Thorac. Surg. 77, 1740–1744 (2004).

11. Solfisburg, Q. S. et al. The Salivary Microbiome and Predicted Metabolite Production Are Associated with Barrett’s Esophagus and High-Grade Dysplasia or Adenocarcinoma. Cancer Epidemiol. Biomarkers Prev. 33, 371–380 (2024).

12. Segal, I., Reinach, S. G. & de Beer, M. Factors associated with oesophageal cancer in Soweto, South Africa. Br. J. Cancer 58, 681–686 (1988).

13. Simba, H., Kuivaniemi, H., Abnet, C. C., Tromp, G. & Sewram, V. Environmental and life-style risk factors for esophageal squamous cell carcinoma in Africa: a systematic review and meta-analysis. BMC Public Health 23, 1782 (2023).

14. Yu, Y. et al. A cross-cohort study identifies potential oral microbial markers for esophageal squamous cell carcinoma. iScience 27, 111453 (2024).

15. Wang, Q. et al. Oral Microbiome in Patients with Oesophageal Squamous Cell Carcinoma. Sci. Rep. 9, 1–9 (2019).

16. Jiang, Z., Wang, J., Qian, X., Zhang, Z. & Wang, S. Oral microbiota may predict the presence of esophageal squamous cell carcinoma. J. Cancer Res. Clin. Oncol. 149, 4731–4739 (2023).

17. Jiang, Z., Wang, J., Shen, Z., Zhang, Z. & Wang, S. Characterization of esophageal Microbiota in patients with esophagitis and esophageal squamous cell carcinoma. Front. Cell. Infect. Microbiol. 11, 774330 (2021).

18. Zhao, Q. et al. Alterations of oral Microbiota in Chinese patients with esophageal cancer. Front. Cell. Infect. Microbiol. 10, 541144 (2020).

19. Zhu, H. et al. Convergent dysbiosis of upper aerodigestive microbiota between patients with esophageal and oral cavity squamous cell carcinoma. Int. J. Cancer 152, 1903–1915 (2023).

20. Annavajhala, M. K. et al. Relationship of the Esophageal Microbiome and Tissue Gene Expression and Links to the Oral Microbiome: A Randomized Clinical Trial. Clin. Transl. Gastroenterol. 11, e00235 (2020).

21. Belstrøm, D. et al. Temporal stability of the salivary Microbiota in oral health. PLoS One 11, e0147472 (2016).

22. Tamashiro, R. et al. Stability of healthy subgingival microbiome across space and time. Sci. Rep. 11, 23987 (2021).

23. Nomoto, D. et al. Fusobacterium nucleatum promotes esophageal squamous cell carcinoma progression via the NOD1/RIPK2/NF-κB pathway. Cancer Lett. 530, 59–67 (2022).

24. Li, Z. et al. Fusobacterium nucleatum predicts a high risk of metastasis for esophageal squamous cell carcinoma. BMC Microbiol. 21, 301 (2021).

25. Whalen, S., Schreiber, J., Noble, W. S. & Pollard, K. S. Navigating the pitfalls of applying machine learning in genomics. Nat. Rev. Genet. 23, 169–181 (2022).

26. Austin, G. I. & Korem, T. Planning and analyzing a low-biomass microbiome study: A data analysis perspective. J. Infect. Dis. jiae378 (2024).

27. Chen, H. et al. Characteristics of the oral and gastric microbiome in patients with early-stage intramucosal esophageal squamous cell carcinoma. BMC Microbiol. 24, 88 (2024).

28. Li, H. et al. Characteristics of oral Microbiota in patients with esophageal cancer in China. Biomed Res. Int. 2021, 2259093 (2021).

29. Li, Z. et al. Characterization of the oral and esophageal Microbiota in esophageal precancerous lesions and squamous cell carcinoma. Front. Cell. Infect. Microbiol. 11, 714162 (2021).

30. Wei, J. et al. Salivary microbiota may predict the presence of esophageal squamous cell carcinoma. Genes & Diseases 9, 1143–1151 (2022).

31. Li, M. et al. Signatures within esophageal microbiota with progression of esophageal squamous cell carcinoma. Chin. J. Cancer Res. 32, 755–767 (2020).

32. Chesdachai, S. et al. The characteristics of Capnocytophaga infection: 10 years of experience. Open Forum Infect. Dis. 8, ofab175 (2021).

33. Zhu, W. et al. Capnocytophaga gingivalis is a potential tumor promotor in oral cancer. Oral Dis. 30, 353–362 (2024).

34. Hawkes, C. G. et al. Selenomonas sputigena interactions with gingival epithelial cells that promote inflammation. Infect. Immun. 91, e0031922 (2023).

35. Cho, H. et al. Selenomonas sputigena acts as a pathobiont mediating spatial structure and biofilm virulence in early childhood caries. Nat. Commun. 14, 2919 (2023).

36. Sasaki, H. et al. Presence of Streptococcus anginosus DNA in esophageal cancer, dysplasia of esophagus, and gastric cancer. Cancer Res. 58, 2991–2995 (1998).

37. Senthil Kumar, S., Johnson, M. D. L. & Wilson, J. E. Insights into the enigma of oral streptococci in carcinogenesis. Microbiol. Mol. Biol. Rev. 88, e0009523 (2024).

38. Fu, K. et al. Streptococcus anginosus promotes gastric inflammation, atrophy, and tumorigenesis in mice. Cell 187, 882–896.e17 (2024).

39. Nomburg, J. et al. An international report on bacterial communities in esophageal squamous cell carcinoma. Int. J. Cancer 151, 1947–1959 (2022).

40. Manson McGuire, A., et al. Evolution of invasion in a diverse set of Fusobacterium species. MBio 5, e01864 (2014).

41. Rubinstein, M. R. et al. Fusobacterium nucleatum promotes colorectal carcinogenesis by modulating E-cadherin/β-catenin signaling via its FadA adhesin. Cell Host Microbe 14, 195–206 (2013).

42. Fardini, Y. et al. Fusobacterium nucleatum adhesin FadA binds vascular endothelial cadherin and alters endothelial integrity. Mol. Microbiol. 82, 1468–1480 (2011).

43. Abnet, C. C. et al. Tooth loss and lack of regular oral hygiene are associated with higher risk of esophageal squamous cell carcinoma. Cancer Epidemiol. Biomarkers Prev. 17, 3062–3068 (2008).

44. Chen, X. et al. Poor oral health is associated with an increased risk of esophageal squamous cell carcinoma - a population-based case-control study in China: Oral health and risk of esophageal squamous cell carcinoma. Int. J. Cancer 140, 626–635 (2017).

45. Dar, N. A. et al. Poor oral hygiene and risk of esophageal squamous cell carcinoma in Kashmir. Br. J. Cancer 109, 1367–1372 (2013).

46. Kaimila, B. et al. Poor oral health and the risk of esophageal squamous cell carcinoma in Malawi. Int. J. Cancer 154, 1587–1595 (2024).

47. Socransky, S. S., Haffajee, A. D., Cugini, M. A., Smith, C. & Kent, R. L., Jr. Microbial complexes in subgingival plaque. J. Clin. Periodontol. 25, 134–144 (1998).

48. Ruan, X., Luo, J., Zhang, P. & Howell, K. The salivary microbiome shows a high prevalence of core bacterial members yet variability across human populations. NPJ Biofilms Microbiomes 8, 85 (2022).

49. Sinha, R. et al. Assessment of variation in microbial community amplicon sequencing by the Microbiome Quality Control (MBQC) project consortium. Nat. Biotechnol. 35, 1077–1086 (2017).

50. Kennedy, N. A. et al. The impact of different DNA extraction kits and laboratories upon the assessment of human gut microbiota composition by 16S rRNA gene sequencing. PLoS One 9, e88982 (2014).

51. Clausen, D. S. & Willis, A. D. Evaluating replicability in microbiome data. Biostatistics 23, 1099–1114 (2022).

52. Langmead, B. & Salzberg, S. L. Fast gapped-read alignment with Bowtie 2. Nat. Methods 9, 357–359 (2012).

53. Bolyen, E. et al. Reproducible, interactive, scalable and extensible microbiome data science using QIIME 2. Nat. Biotechnol. 37, 852–857 (2019).

54. Callahan, B. J. et al. DADA2: High-resolution sample inference from Illumina amplicon data. Nat. Methods 13, 581–583 (2016).

55. Katoh, K., Misawa, K., Kuma, K.-I. & Miyata, T. MAFFT: a novel method for rapid multiple sequence alignment based on fast Fourier transform. Nucleic Acids Res. 30, 3059–3066 (2002).

56. Price, M. N., Dehal, P. S. & Arkin, A. P. FastTree 2--approximately maximum-likelihood trees for large alignments. PLoS One 5, e9490 (2010).

57. Bokulich, N. A. et al. Optimizing taxonomic classification of marker-gene amplicon sequences with QIIME 2’s q2-feature-classifier plugin. Microbiome 6, 90 (2018).

58. Chen, T. et al. The Human Oral Microbiome Database: a web accessible resource for investigating oral microbe taxonomic and genomic information. Database 2010, baq013 (2010).

59. Austin, G. I. et al. Contamination source modeling with SCRuB improves cancer phenotype prediction from microbiome data. Nat. Biotechnol. 41, 1820–1828 (2023).

60. Lozupone, C., Lladser, M. E., Knights, D., Stombaugh, J. & Knight, R. UniFrac: an effective distance metric for microbial community comparison. ISME J. 5, 169–172 (2011).

61. Anderson, M. J. A new method for non-parametric multivariate analysis of variance. Austral Ecol. 26, 32–46 (2001).

62. Martin, B. D., Witten, D. & Willis, A. D. MODELING MICROBIAL ABUNDANCES AND DYSBIOSIS WITH BETA-BINOMIAL REGRESSION. Ann. Appl. Stat. 14, 94–115 (2020).

63. Benjamini, Y. & Hochberg, Y. Controlling the False Discovery Rate: A Practical and Powerful Approach to Multiple Testing. J. R. Stat. Soc. Series B Stat. Methodol. 57, 289–300 (1995).

